# Latent Plasticity of Effector-like Exhausted CD8 T cells contributes to memory responses

**DOI:** 10.1101/2020.02.22.960278

**Authors:** Saravanan Raju, Yu Xia, Bence Daniel, Kathryn E. Yost, Elliot Bradshaw, Elena Tonc, Daniel J. Verbaro, Ansuman T. Satpathy, Takeshi Egawa

**Affiliations:** Department of Pathology and Immunology, Washington University School of Medicine, Saint Louis, MO 63110, USA; Department of Pathology, Stanford University School of Medicine, Stanford, CA 94305, USA; Parker Institute for Cancer Immunotherapy, Stanford University School of Medicine, Stanford, CA 94305, USA

## Abstract

Persistent antigen induces a dysfunctional CD8 T cell state known as T cell “exhaustion” characterized by expression of PD-1 and decreased effector functions. Nevertheless, dysfunctional CD8 T cells can mediate control of antigen burden which is long-lasting. While heterogeneity of exhausted CD8 T cells has been described, the cells which actively proliferate and exert viral control have remained elusive. Here, we define subsets of PD-1^+^ CD8 T cells during chronic infection marked by expression of CX3CR1 with substantial *in situ* proliferation and high expression of granzyme B. Moreover, these cells maintain the effector pool through self-renewal independently of previously defined stem-like cells. Unexpectedly, CX3CR1^+^ CD8 T cells retain plasticity to be reprogrammed to memory cells through expression of TCF-1 and re-gain polyfunctionality. Thus, we define a subset of effector-like exhausted CD8 T cells with capacity to contribute to the memory pool, offering a prime target for novel immunotherapies.

## Introduction

Upon encountering antigen, naive CD8 T cells undergo robust clonal expansion and acquire effector functions for host protection against pathogens and tumors. However, in the context of persistent antigen exposure seen in chronic viral infections, they are subject to an alternative differentiation pathway known as T cell exhaustion, which is characterized by continued expression of inhibitory receptors, decreased cytokine secretion, and blunted proliferation (*1, 2*). Although the acquisition of this dysfunctional state serves as a protective host mechanism by which CD8 T cell-mediated tissue damage is restricted, this pathway facilitates persistence of infection or tumors (*3–5*). Nevertheless, exhausted CD8 T cells are not inert and can actively control antigen burden (*6*). Recent data have highlighted heterogeneity within exhausted CD8 T cells both at the population (*7–10*) and single-cell level (*11–13*). However, identification of a subset undergoing active proliferation with effector gene expression that directly impacts viral control is lacking. Thus, there is considerable interest in defining subpopulations of exhausted CD8 T cells and the developmental relationship between them, as such knowledge provides biomarkers for productive CD8 T cell responses as well as direct targets for future immunotherapies.

Our current understanding of exhausted CD8 T cells is based largely upon division of cells into two major subpopulations in viral infection and tumor models; cells marked with expression of the transcription factor TCF-1 (CXCR5^+^) with stem-like characteristics and TCF-1^−/lo^ Blimp-1^+^ TIM3^+^ cells marked by a more severe exhaustion signature despite expression of effector genes such as granzyme B (*Gzmb*) (*8–10*). The TCF-1^+^ population comprises stem-like CD8 T cells that exhibit substantial proliferative capacity upon PD-1 blockade and is thus thought to continuously replenish the TIM3 cell pool during chronic infection or in tumors.

However, these stem-like cells are largely quiescent under physiological conditions and it is thus unclear to what extent they undergo proliferation and actively contribute to replenishment of the effector CD8 T cell pool (*12*). Alternatively, it is possible, as seen in steady-state hematopoiesis (*14–16*), that the antigen-specific CD8 T cell pool is maintained mainly by intermediate precursors that actively self-renew to some extent and also give rise to differentiated effector cells, whereas stem-like cells only rarely transit to the more mature developmental stages. Yet, the identity of the proliferative subpopulation of CD8 T cells and their connection to the two major subpopulations aforementioned, is poorly understood.

In this work, we identified subpopulations of proliferating CD8 T cells that are marked by the expression of CX3CR1 and require CD4 T cell help, IL-21 and T-bet in the chronic LCMV infection model. Analogous CD8 subpopulations were also found in transplanted tumors. CX3CR1^+^ cells emerge in parallel with TCF-1^+^ and TIM3^+^ cells during the subacute phase of antiviral response, and their appearance is independent of the TCF-1^+^ population. Tamoxifen-inducible fate-mapping revealed they are maintained by self-renewal, and they contribute little to the pool of terminally exhausted TIM3^+^ CX3CR1^−^ CD8 T cells. Unexpectedly, despite their acquisition of Blimp-1 together with loss of TCF-1 expression during antiviral response, CX3CR1^+^ PD-1^+^ CD8 T cells become memory cells with renewed TCF-1 expression and proliferative capacity upon rechallenge. These results demonstrate early segregation of PD-1^+^ CD8 T cells into distinct lineages analogous to effector memory, central memory and differentiated effectors during acute responses, and latent plasticity of exhausted CD8 T cells for their reprogramming to memory T cells.

## Results

### CX3CR1 defines a unique subset of exhausted non-TCF1 CD8 T cells that actively proliferate

To investigate additional heterogeneity within exhausted CD8 T cells, we performed single cell RNA sequencing (scRNA-seq) of antigen-specific gp33-tetramer binding cells during the chronic phase of LCMV-c13 infection. UMAP projection of this dataset demonstrated 4 clusters of cells including the previously defined TCF-1^+^ and TIM3^+^ subsets (**Fig. 1A, Fig. S1A**). Two additional clusters were identified that were most notable for CX3CR1 expression. One of these CX3CR1 clusters was enriched for expression of genes encoding *Tbx21* and *Zeb2* while expression of *Pdcd1*, *Eomes* and *Tox* was low. The other CX3CR1^+^ cluster was marked with increased expression of inhibitory receptors *Havcr2*. *Lag3* and *Tigit*, with a significant proportion of the cells positive for Mki67, encoding Ki-67 protein, suggesting active proliferation. We thus sought to better characterize these populations during chronic viral infection.

**Fig. 1.**
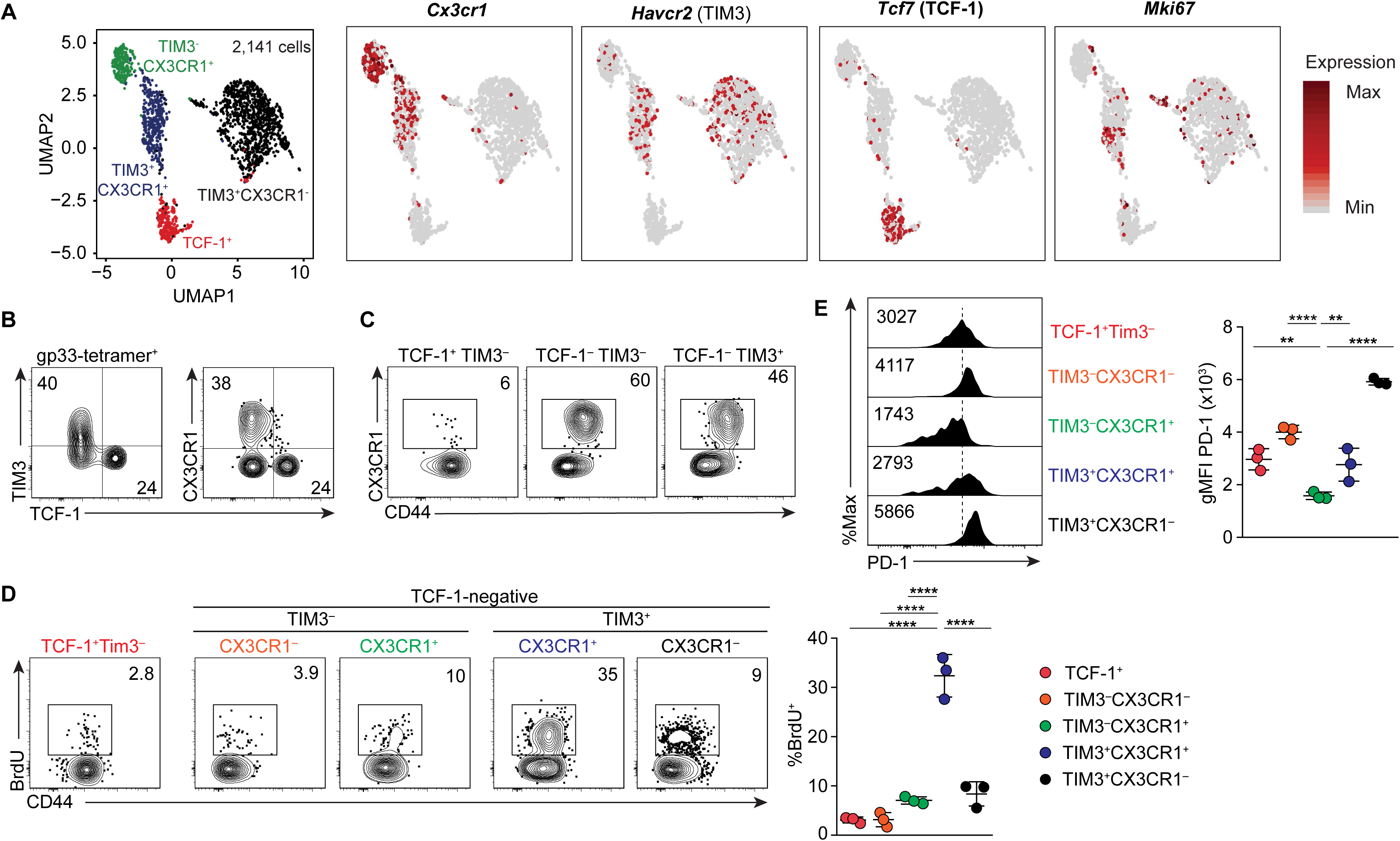
CX3CR1 marks a proliferative subset of exhausted T cells. **A.** UMAP analysis of scRNA-seq data of 2,141 gp-33 specific CD8 T cells on 21 dpi with LCMV-c13. Right panels show the expression of indicated genes. **B, C.** Flow cytometry showing expression of TIM3, CX3CR1, CD44 and TCF-1 in LCMV gp33-specific CD8 T cells 4 weeks after infection of C57BL/6 mice with LCMV-c13. Numbers indicate percentages of gated cells in each parental population shown in the figure. Data are representative of 8 mice. **D.** BrdU incorporation as measured by flow cytometry by LCMV gp33-specific CD8 T cells in the indicated subpopulations 12 hours after intraperitoneal administration of 1 mg BrdU on 22 dpi with LCMV-c13. Data are representative of three independent experiments with n=3 mice each. **E.** Expression of PD-1 measured by flow cytometry in the indicated population of LCMV-gp33-specific CD8 T cells on 22 dpi, quantified in the right panel. Data are representative of 3 independent experiments with n=3 mice each.

Consistent with the scRNA-seq data, approximately 40% of LCMV-gp33-specific CD8 T cells in mice infected with LCMV-clone 13 (LCMV-c13) expressed CX3CR1, and its expression was seen exclusively in TCF-1-negative cells (**Fig. 1B**). The frequency of CX3CR1^+^ cells was notable in TCF-1^−^ TIM3^−^ CD8 T cells, a subpopulation of PD-1^+^ CD8 T cells distinct from the previously characterized TCF-1^+^ and TIM3^+^ cells, and expressed CD44 at higher levels than CX3CR1^−^ cells in both TIM3^−^ and TIM3^+^ fractions (**Fig. 1C**).

To determine whether the CX3CR1^+^ cells comprise proliferating CD8 T cells, we measured BrdU incorporation, which directly reflects DNA synthesis during the S-phase of the cell cycle, following pulse labeling in the chronic phase (22 dpi) of LCMV-c13 infection (**Fig. 1D**). We confirmed the quiescence of TCF-1^+^ cells as only <5% of cells incorporated BrdU over 12-hours following one injection of BrdU. Among the TCF-1^−^ fractions, BrdU^+^ cells were significantly enriched in the CX3CR1^+^ populations with the TIM3^+^ CX3CR1^+^ cells exhibiting by far the highest incorporation of any subset (∼30%). Both TIM3^+^ CX3CR1^−^ and TIM3^−^ CX3CR1^+^ cells incorporated BrdU at lower frequencies than TIM3^+^ CX3CR1^+^ cells while TIM3^−^ CX3CR1^−^ cells were similarly quiescent as TCF-1^+^ stem-like CD8 T cells. Consistent results were seen with Ki-67 staining, which is often used as a surrogate for cell cycle status (**Fig. S1b,c**). CD8 T cells in all the subpopulations expressed Ki-67 at similarly high frequencies during the subacute phase of antiviral response (8 dpi). In the chronic phase (22 dpi), the majority of TIM3^+^ CX3CR1^+^ cells retained Ki-67 expression at a higher level compared to all other populations. These results support the notion that not all TIM3^+^ cells lose proliferative capacity and are terminally exhausted (*12*), and importantly, that expression of CX3CR1 marks proliferating effector cells within TIM3^+^ cells.

The amount of PD-1 surface expression is known to correlate with the extent of exhaustion, with higher levels corresponding to more severe exhaustion and unresponsiveness to the checkpoint blockade (*7, 17, 18*). Prior studies have demonstrated that TCF-1^+^subset expresses lower amounts of PD-1 compared to TIM3^+^ cells, and is thus less exhausted with reserved proliferative potential. Among CD8 T cells in the five subpopulations, TIM3^−^ CX3CR1^+^ cells expressed the lowest amounts of PD-1 compared to all other subsets, including TCF-1^+^ cells, previously shown to express lower amounts of PD-1 compared to TIM3^+^ cells (**Fig. 1E**). TIM3^+^ CX3CR1^−^ cells expressed the highest levels of PD-1, consistent with a more severely exhausted state.

A critical aspect of T cell biology is egress of CD8 T cells from secondary lymphoid organs to peripheral blood and extravasation into tissue to mediate immunity. Effector cell subsets are typically found in both blood and tissue at higher frequencies compared to memory-like cells (*19*). To examine the migratory behavior of CX3CR1^+^ cells, we performed *in vivo* labelling of T cells by an intravenous injection of anti-CD45.2 which allowed us to discriminate between tissue resident (CD45.2^−^) and circulating (CD45.2^+^) antigen-specific CD8 T cells present in lung. In contrast to TIM3^+^ CX3CR1^−^ cells that were predominantly localized in the lung parenchyma, distribution of both CX3CR1^+^ subpopulations was significantly enriched in the circulating compartment (**Fig. S1D**). Within the CX3CR1^+^ populations, more TIM3^+^ cells were found in the tissues compared to TIM3^−^ cells, suggesting that TIM3^+^ CX3CR1^+^ cells preferentially migrate to the tissue, and proliferate upon antigen recognition.

We then asked whether analogous CX3CR1^+^ PD-1^+^ CD8 T cells could be found in tumor-infiltrating lymphocytes (TILs) which are also exposed to chronic antigen stimulation. Indeed, when we analyzed TILs in MC38 and E.G7 tumors following subcutaneous inoculation we observed a fraction of PD-1^+^ cells that expressed CX3CR1 (**Fig. S2A,B**). These results highlight unique characteristics of CX3CR1^+^ PD-1^+^ CD8 T cells, which are distinct from the previously characterized exhausted CD8 T cells in their proliferative activity and tissue distribution, and are present in both chronic infection and tumors.

### CX3CR1^+^ PD-1^+^ CD8 T cells comprise cells with active cell cycling and expression of effector molecules

While scRNA-seq analysis allows characterization of novel cell populations, full transcriptome analysis is technically challenging due to low sequencing depth. To gain additional insight into the CX3CR1^+^ populations of CD8 T cells, we purified two subsets of CX3CR1^+^ cells based on TIM3 expression as well as stem-like (TCF-1^+^) and terminally exhausted (TIM3^+^ CX3CR1^−^) CD8 T cells for bulk RNA-seq on day 16 of LCMV infection. Unsupervised clustering and principal component analysis showed that CX3CR1^+^ populations are transcriptomically distinct from TCF-1^+^ stem-like cells and TIM3^+^ CX3CR1^−^ CD8 T cells (**Fig. 2A** and **Fig. S3**). Among genes encoding transcription factors, CX3CR1^+^ populations expressed high *Tbx21* and *Zeb2,* which cooperates with T-bet to regulate effector function of CD8 T cells as well as *Prdm1* encoding BLIMP-1(*20–22*) (**fig. 2B**). A few factors that have been associated with memory CD8 T cells and stem-like PD-1^+^ CD8 T cells, including *Tcf7*, *Id3* and *Bcl6*, were expressed at substantially lower levels by the CX3CR1^+^ cells compared to the TCF-1^+^ stem-like population (**fig. 2B**). Expression of genes associated with CD8 effector functions was also distinct between TCF-1^+^ stem-like and CX3CR1^+^ populations. Consistent with elevated *Tbx21* and *Prdm1*, expression of *Ifng*, *Prf1*, *Gzma* and *Gzmb* was higher in CX3CR1^+^ populations compared to TCF-1^+^ cells although these genes were more highly expressed in TIM3^+^ than TIM3^−^ populations at the population level (**fig. 2B**). Gene Set Enrichment Analysis (GSEA) also showed significant enrichment of genes associated with pathways for cell proliferation and T cell effector function in TIM3^+^ CX3CR1^+^ cells compared to TCF-1^+^ stem-like cells (**fig. 2C**). These results suggest that TIM3^+^ CX3CR1^+^ cells are the most activated and proliferative effectors among the antigen-specific PD-1^+^ CD8 T cells.

**Fig. 2.**
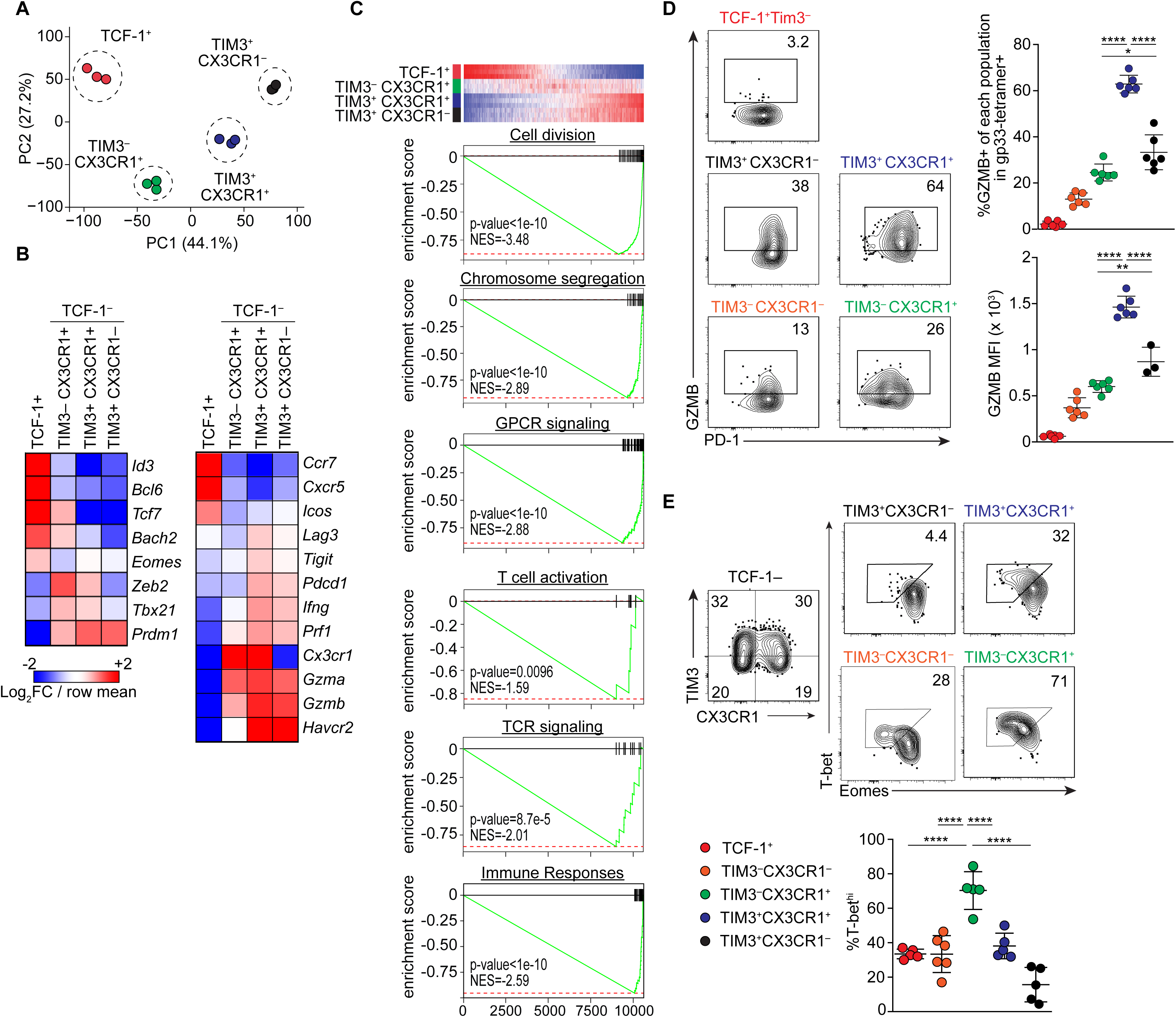
Gene expression analysis of exhausted CD8 T cell subsets defined by CX3CR1. **A.** A Principal Component Analysis (PCA) plot derived from bulk RNA-seq analysis of the indicated subpopulations of PD-1^+^ CD8 T cells on 16 dpi. **B.** Heat maps showing expression of indicated genes by distinct subsets of PD-1^+^ CD8 T cells. Expression values were averaged from three independent samples for each population and color-coded based on row mean and fold-changes. **C.** Gene Set Enrichment Analysis (GSEA) showing pathways associated with genes that were differentially expressed between TCF-1^+^ and TCF-1^−^ TIM3^+^ CX3CR1^+^ populations. **D.** Expression of Granzyme B (GZMB) protein by gp33-specific CD8 T cells shown by frequencies of cells expressing GZMB and by mean fluorescence intensity (MFI) on 28 dpi. Data are representative of 2 independent experiments with n>5 mice in total. **E.** Flow Cytometry plots showing expression of T-bet and Eomes in the indicated subsets of gp33-specific CD8 T cells in the spleen on 22 dpi with LCMV-c13 infection. Data are representative of 3 independent experiments with n=5 mice each. Dots in the graphs in (A), (D) and (E) indicate individual mouse and data in (D) and (E) are shown by mean ± SD. Statistical differences in (D) and (E) were tested using one-way ANOVA with a Tukey post-hoc test.

We further examined amounts of proteins associated with CD8 T cell effector function. Consistent with the gene expression profiling data, expression of granzyme B (GZMB) by LCMV-specific CD8 T cells *ex vivo* was highest in the TIM3^+^ CX3CR1^+^ cells among the PD-1^+^ CD8 T cells (**fig. 2D**). Despite comparable *Gzmb* mRNA expression between TIM3^+^ CX3CR1^+^ cells and TIM3^+^ CX3CR1^−^ cells, GZMB protein was expressed at a higher frequency and level in TIM3^+^ CX3CR1^+^ cells, further suggesting that they retain effector function as cytotoxic T cells. Furthermore, we profiled expression of the two E-box transcription factors T-bet and Eomes, which are partially redundant in the control of effector functions of CD8 T cells (*23*). While a previous study demonstrated that T-bet is necessary for the development of PD-1^lo/int^ progenitor-like CD8 T cells, the nature of the T-bet^hi^ Eomes^lo^ PD-1^lo/in^ progenitor-like cells remains unknown since TCF-1^+^ stem-like CD8 T cells are Eomes^hi^ (*8*). Among the four TCF-1^−^ subpopulations in the LCMV-specific CD8 T cells, TIM3^−^ CX3CR1^+^ cells expressed significantly higher levels of T-bet followed by TIM3^+^ CX3CR1^+^ cells (**fig. 2E**). In contrast, TIM3^+^ CX3CR1^−^ cells were devoid of T-bet-expressing cells. These results indicate that CX3CR1^+^ cells are enriched for cells with active effector functions, while TIM3^+^ CX3CR1^−^ cells are terminally exhausted.

### CD4 help mediated by IL-21 is required for exhausted CX3CR1^+^ cells

We next sought to determine what factors within the immune response promote the generation and/or maintenance of the CX3CR1^+^ populations. It is well established that the absence of CD4 T cells in LCMV-c13 infection causes severe CD8 T cell exhaustion, allowing permanent persistence of virus (*2*). Severe CD8 T cell exhaustion was observed early in infection, when viral burden is still comparable between CD4 T cell-replete and -depleted conditions (*7, 24*). Considering this, we asked whether the progressive CD8 T cell exhaustion in the “helpless” context results from the defective development of CX3CR1^+^ CD8 T cell populations. Indeed, in mice treated with anti-CD4 prior to infection with LCMV-c13, we observed an almost complete absence of CX3CR1^+^ cells in LCMV-specific, PD-1^+^ CD8 T cells 3-4 weeks after infection (**fig. 3A,B, fig. S4**). Consistent with the enrichment of T-bet^+^ cells in the CX3CR1^+^ fractions, overall T-bet^+^ cells in LCMV-gp33-specific CD8 T cells were severely reduced in CD4-depleted mice (**fig. 3c**). Analysis of CD4-depleted versus CD4-replete, LCMV-c13-infected mice in the subacute period on 8 dpi showed relatively normal development of LCMV-specific CX3CR1^+^ CD8 T cells, including TIM3^+^ and TIM3^−^ subpopulations (**fig. 3A,B**). On 15 dpi however, TIM3^−^ CX3CR1^+^ CD8 T cells were substantially reduced while TIM3^+^ CX3CR1^+^ CD8 T cells were less severely affected, followed by almost total loss of CX3CR1^+^ CD8 T cells on 22 dpi. Despite the drastic reduction of CX3CR1^+^ cells, the numbers of TIM3^+^ CX3CR1^−^ cells were comparable between CD4-depleted and CD4-replete mice (**fig. 3B**). These results indicate that the presence of CD4 T cells is dispensable for the initial development but necessary for the maintenance of TIM3^−^ CX3CR1^+^ T-bet^Hi^ cells, which may function as the precursor for TIM3^+^ CX3CR1^+^ cells. Our results also suggest that the TIM3^+^ CX3CR1^−^ cells are maintained independently of CX3CR1^+^ cells.

**Fig.3.**
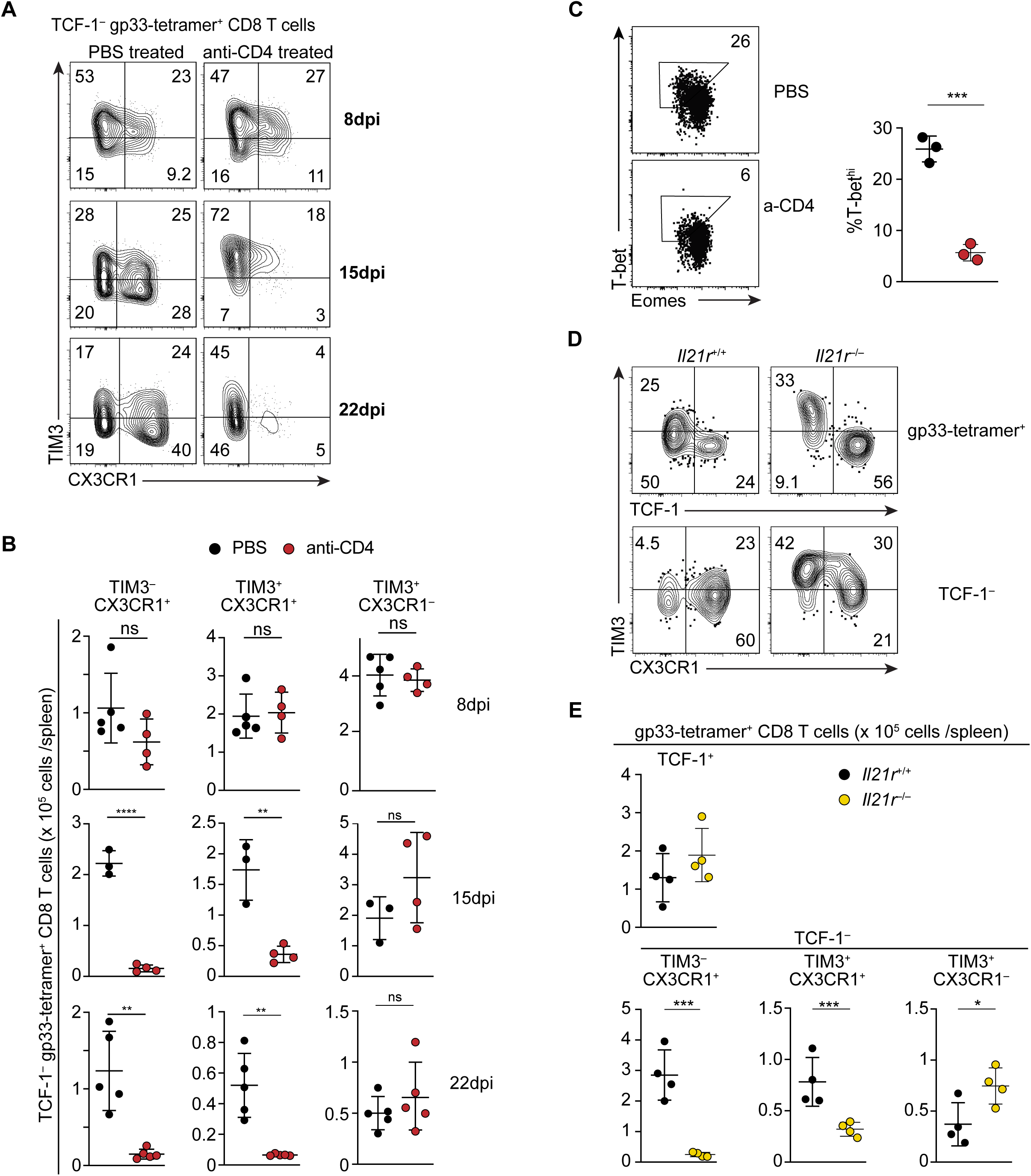
IL-21 and CD4 T cells are required for the maintenance of CX3CR1 subset of PD-1+ CD8 T cells. **A, B.** Flow Cytometry showing expression of CX3CR1 and TIM3 (A) in TCF-1^−^ gp33-specific CD8 T cells in control and CD4 T cell-depleted mice on the indicated dpi with LCMV-c13. (A) shows representative plots from 4-5 mice and data from replicates are shown in (B). **C.** Expression of T-bet and Eomes in in gp33-specific CD8 T cells in control and CD4 T cell-depleted mice on 30 dpi. Frequencies of T-bet^hi^ Eomes^lo^ cells are indicated with polygon gates. Data are representative from 3 independent experiments with n=3 mice in each. **D, E.** Flow Cytometry showing expression of TCF-1, CX3CR1, and TIM3 in gp33-specific cells in spleen of *Il21r*^+/+^ and *Il21r*^−/–^ mice on 35 dpi with LCMV-c13 (D). Data are representative of two independent experiments with n>3 mice in each experiment per genotype. Data in (B), (C), and (E) are shown with mean ± SD.

CD4 T cells enhance CD8 T cell responses by providing direct help to CD8 T cells through cytokine secretion and licensing of dendritic cells. Among molecules involved in CD4 T cell-mediated help, IL-21 is predominantly produced by CD4 T cells and required to sustain CD8 T cell responses to chronic LCMV infection (*25–27*). We thus determined whether IL-21 is necessary for the CD4 T cell-dependent maintenance of the CX3CR1^+^ populations of PD-1^+^ CD8 T cells. On 35 dpi, we observed a significant reduction in CX3CR1-expressing PD-1^+^ CD8 T cells in *Il21r*^−/–^ mice (**fig. 3D, E, fig. S5A,B**). Consistent with CD4 T cell-independent development (*8*), the absolute numbers of TCF-1^+^ stem-like CD8 T cells were comparable between *Il21r*^+/+^ and *Il21r*^−/–^ mice. However, TCF-1^−^ PD-1^+^ CD8 T cells were reduced by 3-fold and this reduction was predominantly caused by the profound loss of CX3CR1^+^ cells (**fig. 3D,E**). These results suggest that CD4 T cell-derived IL-21 enhances the development or maintenance of TIM3^−^ CX3CR1^+^ cells, possibly through the induction of T-bet expression (*28*). Similarly to CD4 depleted mice, numbers of TIM3^+^ CX3CR1^−^ cells were comparable between *Il21r*^+/+^ and *Il21r*^−/–^ mice (**fig. 3E**), further suggesting that the maintenance of TIM3^+^ CX3CR1^−^ cells is independent of CX3CR1^+^ populations. Although follicular helper T (Tfh) cells in germinal centers produce IL-21, we observed intact CX3CR1^+^ PD-1^+^ CD8 T cells in muMT (*Ighm*^−/–^) mice lacking B cells (**fig. S5C,D**), suggesting that the production of IL-21 by CD4 T cells does not require TFH differentiation in the germinal center.

### CX3CR1^+^ expressing CD8 T cells are uniquely dependent on T-bet

Our gene expression profile highlighted a reciprocal expression of the two E-box proteins, T-bet and Eomes in distinct subpopulations of antigen-specific PD-1^+^ CD8 T cells. To determine how dichotomous expression of T-bet and Eomes imprints a molecular program onto exhausted CD8 T cells, we infected mice in which *Tbx21* or *Eomes* was deleted in CD8 T cells. During the chronic phase of LCMV infection (29 dpi), frequencies of total LCMV-specific CD8 T cells were reduced by 2.5-fold in CD8-cre *Tbx21*^F/F^ mice (**fig. 4A,B**). This reduction of total antigen-specific CD8 T cells was attributed to marked reduction of CX3CR1^+^ CD8 T cells, most notably to TIM3^−^ CX3CR1^+^ that expressed the highest amounts of T-bet among all fractions (**Figs. 2E, 4A,B**). Notably, the reduction of TCF-1^+^ stem-like cells as well as TIM3^+^ CX3CR1^−^ cells was minimal, indicating that T-bet plays the crucial role in specifically programming the CX3CR1^+^ subpopulations of exhausted CD8 T cells (**fig. 4A,B**). To determine the impact of loss of CX3CR1^+^ CD8 T cells on viral control, we tracked LCMV-c13 infected CD8-cre *Tbx21*^F/F^ and control *Tbx21*^F/F^ mice for 100 days, by which time the majority of immune-competent mice resolve viremia (**fig. 4C**). While WT mice were able to resolve viremia, T-bet-deficient mice still demonstrated significant viremia. These results indicate deletion of T-bet/*Tbx21* specifically results in loss of CX3CR1^+^ effector cells required for viral control. Importantly, while only representing a fraction of total antigen-specific PD-1^+^ CD8 T cells, this population is crucial for an overall productive CD8 T cell response.

**Fig.4.**
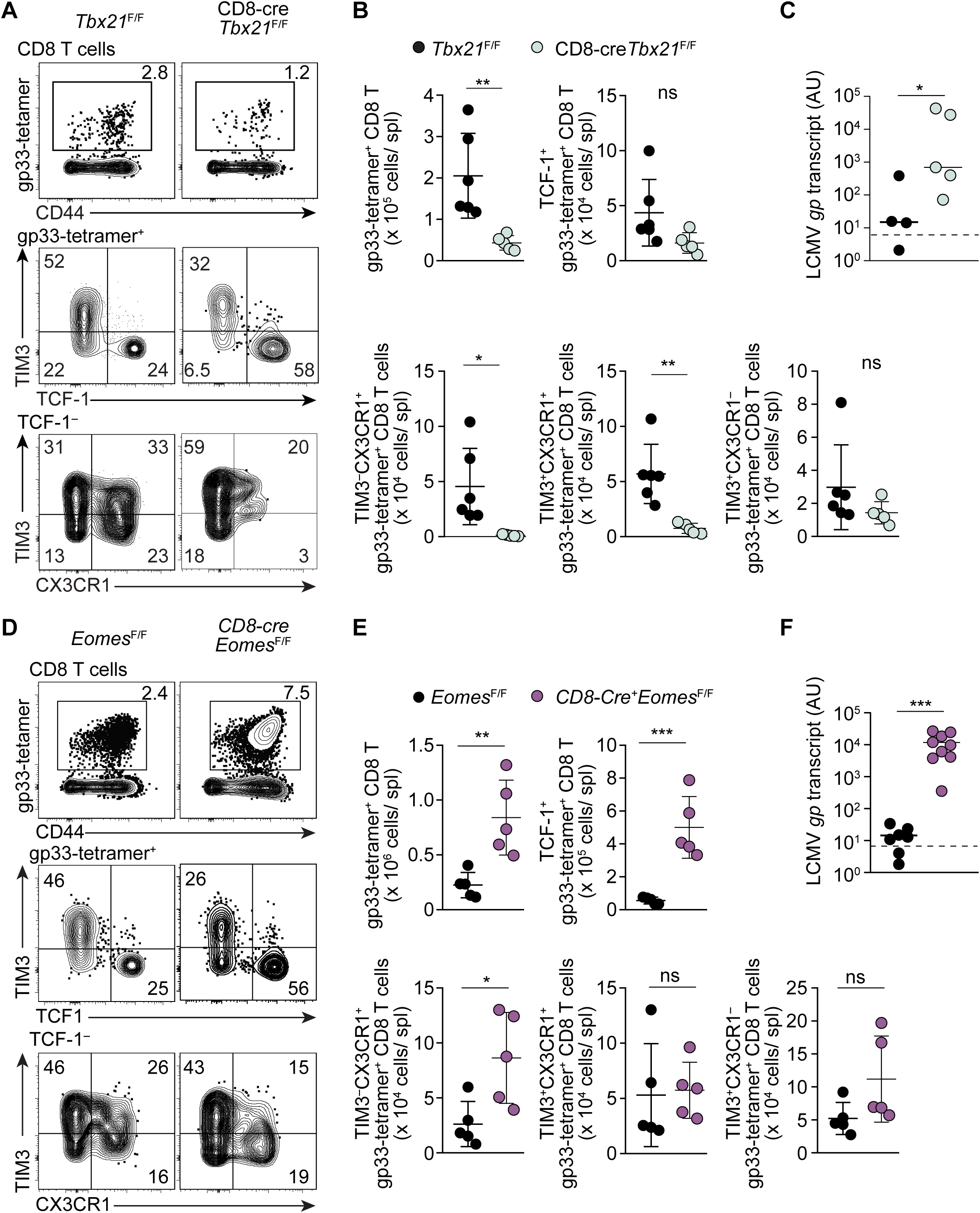
T-bet and Eomes are non-redundantly required for differentiation of PD-1^+^ CD8 T cells in response to chronic viral infection. **A, B.** Flow Cytometry showing expression of TCF-1, CX3CR1, and TIM3 by gp33-specific CD8 T cells in control *Tbx21*^F/F^ and CD8-cre *Tbx21* ^F/F^ mice on 29 dpi with LCMV-c13. Representative flow cytometry plots are shown in (A) with frequencies of gated cells in each parental population, and statistical analyses are shown in (B). Data are representative of 3 independent experiments with n>4 mice per genotype in each experiment. **C.** LCMV-gp mRNA abundance in plasma as assessed by qRT-PCR in *Tbx21*^F/F^ and CD8-cre *Tbx21*^F/F^ mice on 100 dpi with LCMV-c13. Horizontal bars indicate median for samples in each genotype and the statistical differences were assessed by Mann-Whitney U-test. **D, E.** Flow Cytometry showing expression of TCF-1, CX3CR1, and TIM3 in control *Eomes*^F/F^ and CD8-cre *Eomes*^F/F^ mice on 28 dpi with LCMV-c13. Data are representative of 3 independent experiments with n>4 mice per genotype in each experiment. **F.** Viral Load as assessed by qRT-PCR in plasma from *Eomes*^F/F^ and CD8-cre *Eomes*^F/F^ mice on 100 dpi with after LCMV-c13. Data in (B) and (E) are shown with mean ± SD with statistical tests using unpaired *t*-test.

In contrast to T-bet expression being highest in TIM3^−^ CX3CR1^+^ cells, Eomes was expressed more broadly in PD-1^+^ CD8 T cells with its expression lowest in TIM3^−^ CX3CR1^+^ cells (**fig. 2E**). Deletion of *Eomes* in CD8 T cells resulted in increased frequencies and absolute numbers of total LCMV-gp33-specific CD8 T cells approximately by 3-fold (**fig. 4D, E**). This increase was most attributable to a marked increase in TCF-1^+^ stem-like CD8 T cells (**fig. 4E**). In the TCF-1^−^ fractions, however, we did not detect significant differences except for a ∼2.5-fold increase in the numbers of TIM3^−^ CX3CR1^+^ CD8 T cells, suggesting that Eomes functions as a negative regulator for mobilization of relatively quiescent CD8 T cells, including TCF-1^+^ and TIM3^−^ CX3CR1^+^ cells, to downstream proliferation and differentiation programs. Finally, *Eomes* in CD8 T cells was essential for control of chronic LCMV infection with viral persistence in CD8-cre *Eomes*^F/F^ mice with the majority of CD8-cre *Eomes*^F/F^ mice failing to resolve viremia on ≥100 dpi (**fig. 4F**). These results together indicate non-redundant roles of the two E-box proteins, T-bet and Eomes, in CD8 T cell responses to persisting antigen, and a unique role for T-bet in the maintenance of CX3CR1 expressing cells.

### CX3CR1^+^ cells comprise a distinct lineage that develop independently of the TCF-1^+^ stem-like population and are maintained by self-renewal

Adoptive transfer studies demonstrated that TCF-1^+^ stem-like CD8 T cells give rise to terminally differentiated TIM3^+^ cells (*8*). However, it remains unknown whether CX3CR1^+^ T cells are derived from TCF-1^+^ cells as intermediate cells that eventually give rise to terminally differentiated cells, or they constitute a distinct lineage of CD8 cells that develop and is maintained through a unique pathway. To address this question, we first utilized mice in which differentiation of the TCF-1^+^ population is defective. Among the transcription factors selectively expressed in the TCF-1^+^ stem-like CD8 T cells (**fig. 2B**), deletion of *Bcl6* with CD8-cre in LCMV-c13 infection resulted in a substantial loss of the TCF-1^hi^ population with a small fraction of TCF-1^−/lo^ TIM3^−^ cells remaining on 8 dpi (**fig. S6A**). However, expansion of gp33-specific CD8 T cells was minimally affected by the absence of *Bcl6* with the absolute number of CX3CR1^+^ cells as well as total LCMV-gp33-specific CD8 T cells unchanged or only marginally decreased in CD8-cre *Bcl6*^F/F^ mice compared to control *Bcl6*^F/F^ mice (**fig. S6B**). These results indicate that the initial differentiation of CX3CR1^+^ populations is independent of TCF-1^+^ stem-like cells.

To determine whether CX3CR1^+^ CD8 T cells are constitutively replenished by CX3CR1^−^ cells, presumably by TCF-1^+^ stem-like cells, we fate-mapped CX3CR1^+^ cells by utilizing the *Cx3cr1*-creERT2 allele crossed to the *ROSA26*-loxP-stop-loxP-tdTomato (*R26*tdT) reporter. One dose of tamoxifen (TAM) on 8 dpi was sufficient to label approximately 80% of CX3CR1^+^ CD8 T cells, which was examined by detection of td-Tomato (tdT) fluorescence in CX3CR1^+^ CD8 T cells in the peripheral blood two days after TAM administration (**fig.5A, B, fig. S7A**). Labelling was specific to CX3CR1^+^ as minimal labelling of non-CX3CR1 expressing cells was observed following tamoxifen administration (**fig. S7A,B**). After a two-week chase period, we were surprised to see, the frequencies of tdT^+^ cells in the CX3CR1^+^ CD8 T cells remained high with only a modest decay to 60% on 22 dpi (**fig. 5B**). Frequencies of tdT^+^ cells were the highest in the TCF-1^−^ TIM3^−^ CX3CR1^+^ population followed by the TIM3^+^ CX3CR1^+^ population, while the frequency of TIM3^+^ CX3CR1^+^ cells was higher than that of TIM3^−^ CX3CR1^+^ cells reflecting more active proliferation of TIM3^+^ CX3CR1^+^ cells (**fig. 5C,D**). We also observed approximately 25% of tdT^+^ cells showing the TIM3^+^ CX3CR1^−^ phenotype on 22 dpi (**fig. 5C**). These results indicate that the CX3CR1^+^ lineage is specified early after infection with minimal replenishment by CX3CR1^−^ cells and although expression of CX3CR1 may be upregulated promiscuously by a fraction of antigen-specific CD8 T cells, CX3CR1^+^ cells may retain some plasticity to be converted terminally exhausted TIM3^+^ CX3CR1^−^ cells in the subacute phase.

**Fig.5.**
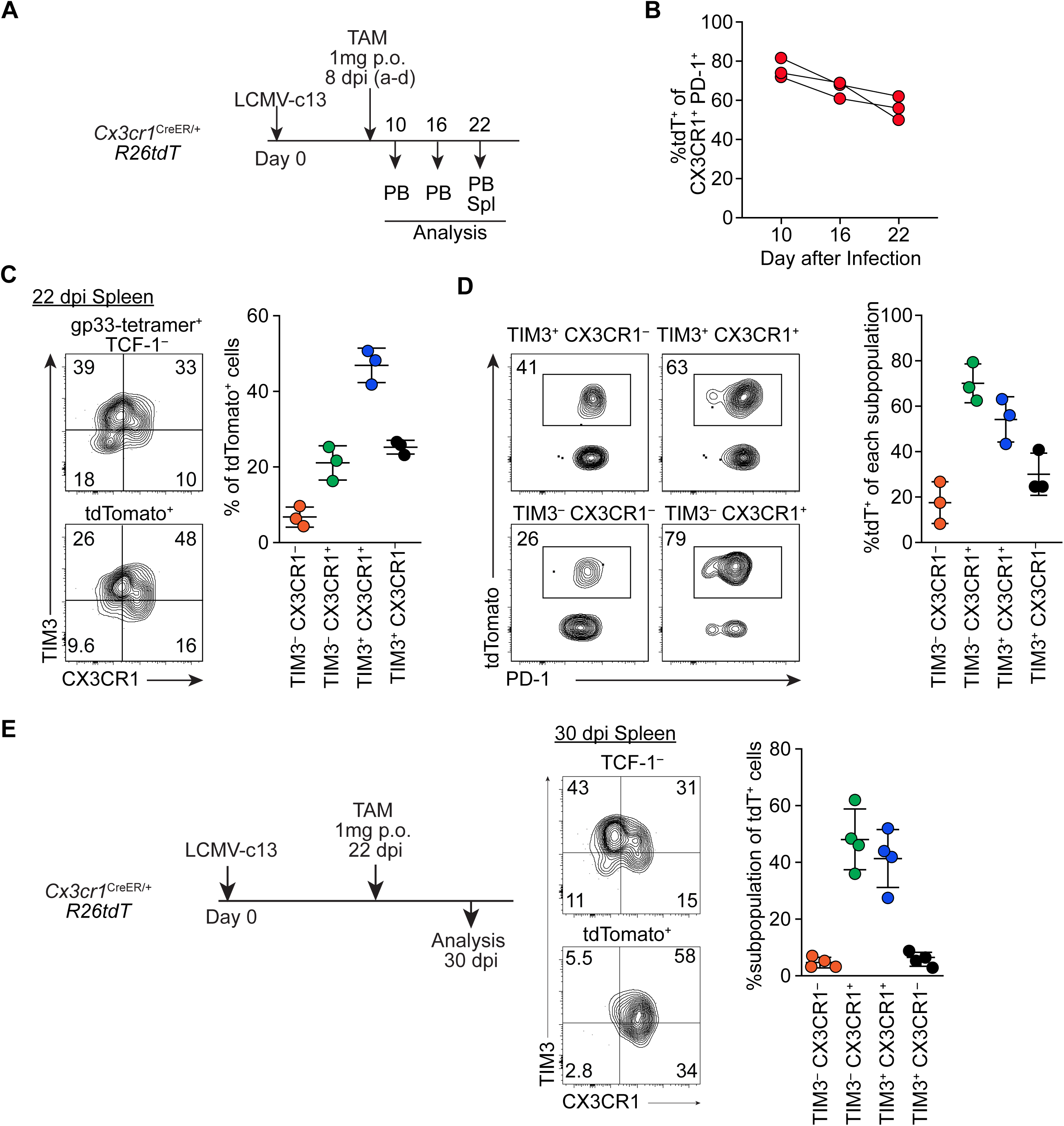
Lineage tracing reveals substantial stability and persistence of CX3CR1^+^ PD-1^+^ CD8 T cells. **A.** Experimental schematic of tamoxifen administration to *Cx3cr1*^creERT2/+^ *ROSA26*^LSL-tdT/+^ (*R26*tdT) mice infected with LCMV-c13 and analysis (B-E). **B.** Frequency of td-Tomato (tdT)^+^ cells within CX3CR1^+^ PD-1^+^ CD8 T cells in the PBMC in *Cx3cr1*^CreER/+^ *R26*tdT mice at indicated time points after infection and tamoxifen administration in (A). Data are representative of 2 independent experiments with at least 3 mice each. **C.** Flow cytometry plots showing expression of CX3CR1and TIM3 in total and tdT^+^ TCF-1^−^ gp33-specific CD8 T cells in the spleen of *Cx3cr1*^creERT2/+^ *R26*tdT mice treated as shown in (A). Pooled data from two independent experiments are shown with mean ± SD. **D.** Frequencies of tdT^+^ cells within indicated subpopulations from samples in (C), quantified in the panel. **E.** Flow cytometry plots showing expression of CX3CR1 and TIM3 in total and tdT^+^ TCF-1^−^ gp33-specific CD8 T cells in the spleen of *Cx3cr1*^creERT2/+^ *R26*tdT mice, which were given Tamoxifen on 22 dpi and analyzed on 30 dpi, quantified in the right panel. Data are representative of 2 independent experiments with n=4 mice each.

When TAM was given in the chronic phase of anti-LCMV response on 22 dpi and labeled cells were chased for 8 days, the vast majority of tdT^+^ cells were found in the CX3CR1^+^ populations (**fig. 5E, fig. S7B,C**). Unlike the early labeling, tdT^+^ cells were excluded from the TCF-1^+^ and TIM3^+^ CX3CR1^−^ populations. Together with the results from the TCF-1^+^ cell-independent formation of CX3CR1^+^ cells in *Bcl6*-CKO mice and their active proliferative activity, these results indicate that CX3CR1^+^ populations constitute a separate lineage which is distinct from TCF-1^+^ and terminally differentiated TIM3^+^ cells and are maintained by self-renewal.

### CX3CR1^+^ cells contribute to long-term memory CD8 T cells expressing TCF-1 after clearance of infection

In immune competent mice, infection of LCMV-c13 is eventually resolved by both T and B cell responses around 100 dpi (*29–31*). We next asked the long-term fate of CX3CR1^+^ cells following the clearance of LCMV-c13. Since it is technically difficult to study memory cell persistence following establishment of T cell exhaustion and antigen clearance in mouse tumor models, use of the LCMV model provides us with a unique opportunity to study long-term fate of CD8 T cells that have gone through exhausting conditions. To this goal, we administered TAM on 28 dpi, which allowed us to minimize the background labeling of TCF-1^+^ CD8 T cells, and chased the tdT-labeled cells for 100-120 days. Similar to acute LCMV infection, we were able to detect the persistence of LCMV-gp33-specific CD8 memory cells in the peripheral blood mononuclear cells and splenocytes of LCMV-c13-immune mice, although they retained low PD-1 expression as reported (*32*) (**fig. 6A**, data not shown). In contrast to LCMV-Arm-immune mice, in which memory CD8 T cells were mostly CD62L^+^ TCF-1^+^ CX3CR1^−^ central memory CD8 T cells, a substantial fraction of gp33-specific memory CD8 T cells in LCMV-c13-immune mice showed the effector-memory phenotype with expression of CX3CR1 (**fig. 6A**). TAM administration to LCMV-c13-infected *Cx3cr1*-creER fate-mapping mice on 28 dpi resulted in marking of ∼40% of LCMV-gp33-specific CD8 memory T cells on 100-120 dpi (**fig. 6A,B**). Almost all tdT^+^ cells continued to express surface CX3CR1 and ∼40% of the CX3CR1^+^ or tdT^+^ cells expressed the memory transcription factor TCF-1 (**fig. 6A,B**). The majority (∼80%) of CX3CR1^+^ memory CD8 T cells were also tdT^+^ and thus were derived from CX3CR1^+^ cells during the labeling period (**fig. 6B**). In contrast to PD-1^+^ CD8 T cells in the chronic phase, the CX3CR1^+^ and CX3CR1^−^ memory CD8 T cells were able to produce IFN-γ following stimulation with LCMV-gp33 or LCMV-gp276 peptide *ex vivo* (**fig. 6C**). Notably, approximately 50% of IFN-g^+^ cells also expressed TNF-a, indicating that the polyfunctionality was restored in these memory CD8 T cell populations (**fig. 6C**). Furthermore, both CX3CR1^+^ and CX3CR1^−^ memory CD8 T cells from LCMV-c13-immune mice were capable of secondary expansion following adoptive transfer to congenic WT mice followed by LCMV-Arm infection (**fig. 6D**). Since TCF-1 was substantially downregulated or not expressed by CX3CR1^+^ cells during the chronic phase in the infected mice, and *Cx3cr1*-creER barely labelled the TCF-1^+^ cells when Tamoxifen was given in the chronic phase, these results indicated that the substantial fraction of TCF-1^+^ memory cells are converted from TCF-1^−^ PD-1^+^ partially exhausted effector cells potentially through de-differentiation. These results collectively indicate that CD8 T cells that fall into the exhausted state with downregulation of memory factors, TCF-1 and BCL6, and upregulation of BLIMP-1 in face of chronic antigen still retain plasticity to acquire the memory phenotype evidenced by long-term persistence and the capability of multiple cytokine production and secondary expansion.

**Fig.6.**
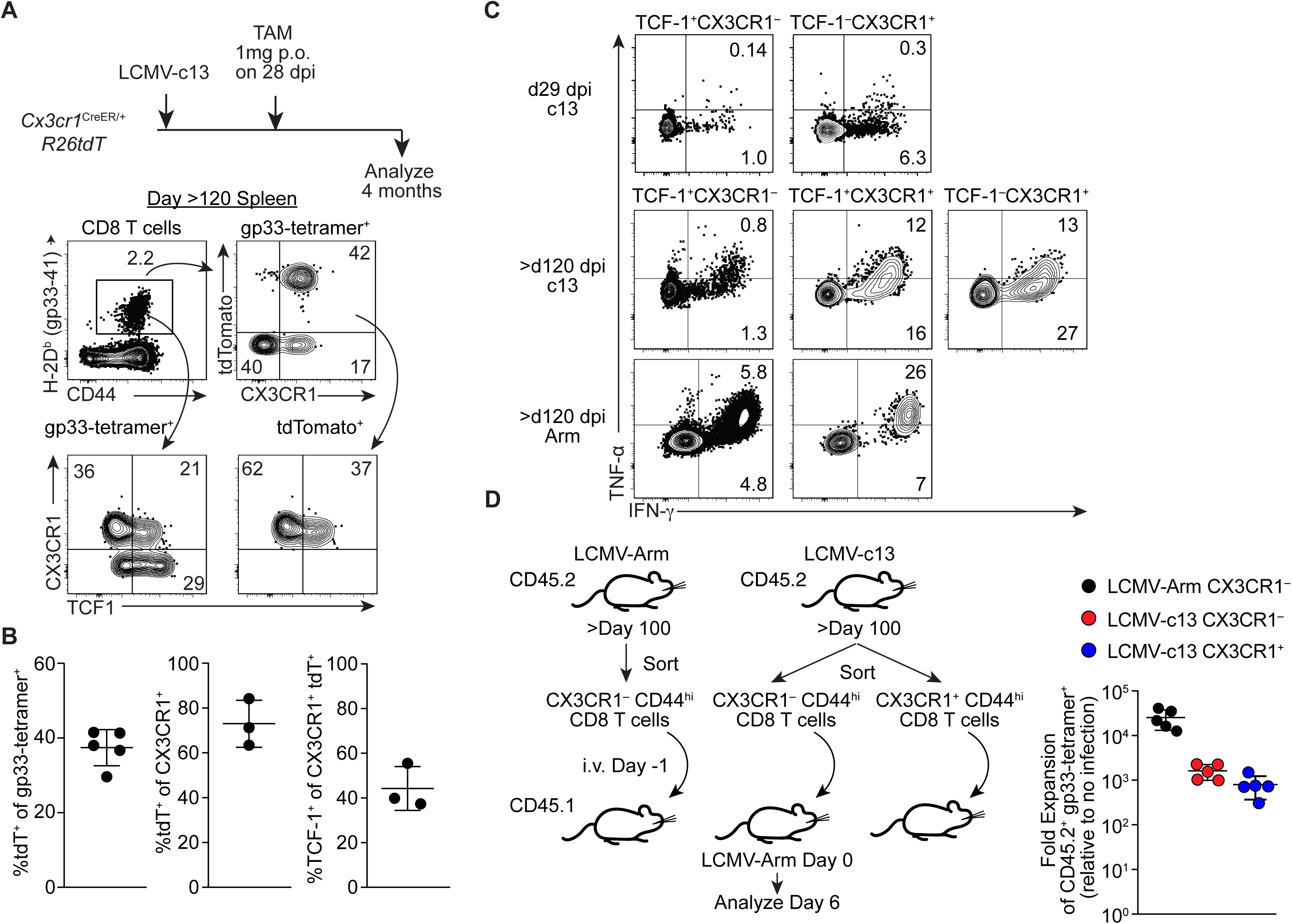
CX3CR1^+^ CD8 T cells during the chronic phase give rise to TCF-1^+^ memory cells following viral clearance. **A, B.** Flow cytometry plots showing expression of tdTomato (tdT), TCF-1, CX3CR1 and TIM3 in gp-33-specific CD8 T cells in the splenocytes in LCMV-c13-infected *Cx3cr1*^creERT2/+^ *R26*tdT mice treated with Tamoxifen on 28 dpi and analyzed on >120 dpi. Representative flow plots from two independent experiments are shown in (A) and data from replicates are shown with mean ± SD in (B). **C.** Expression of IFN-γ and TNF-α in indicated subsets of CD8 T cells in the spleen of d29 dpi LCMV-c13-infected mice, >120 dpi LCMV-Armstrong-infected mice, or >120 dpi LCMV-c13-infected mice following stimulation with LCMV-gp33 peptide *ex vivo* for four hours. **D.** Schematic of adoptive transfer experiment to assess the capability of secondary expansion of CX3CR1^+^ and CX3CR1^−^ CD8 T cells that were sorted from LCMV Arm and LCMV-c13 infected mice. The right panel shows data from replicates. Data are representative from 2 independent experiments with n=2 donor mice and n=5 recipient mice per experiment.

## Discussion

The features of CD8 T cell responses to persistent antigen and mechanistic pathways to control pathogen and tumor burden are of great interest in the development of immunotherapies. CD8 T cells acquire an “exhausted” phenotype in the tumor microenvironment and during response to chronic viral infection (*33*). Recent studies have described heterogeneity of exhausted CD8 T cells in both humans and mouse models, and have identified a TCF-1^+^ stem-like population that retains proliferative potential and responds to immune checkpoint blockade targeting the PD-1/PD-L1 interaction (*8–10*). These findings revealed the heterogeneity of exhausted CD8 T cells associated with their functionality and offer a potential explanation of why only a subset of cancer patients respond to anti-PD-1 therapy. However, subpopulations of exhausted CD8 T cells that are associated with ongoing immune responses have not been defined. Here we identify populations of exhausted CD8 T cells marked by expression of CX3CR1. These cells are endowed with significant proliferative activity, effector functions and capacity to control viral burden. While the CX3CR1^+^ CD8 T cells may be insufficient to eradicate tumors by themselves, the durable CD8 T cell response mediated by these cells can synergize with other immune components to enhance control of virus or tumors. The presence of these cells could thus be a biomarker for patients who respond well to immune checkpoint blockade, as suggested by a report with a small patient cohort (*34*).

Our data showed that the three major populations of PD-1^+^ CD8 T cells, namely TCF-1^+^ stem-like, CX3CR1^+^ proliferative and TIM3^+^ CX3CR1^−^ terminally differentiated/exhausted populations, are segregated early during antiviral responses and maintained independently of one another. This is reminiscent of the separation among terminal effectors, central memory and effector memory cells, which are also marked by CX3CR1 expression during acute responses, suggesting that analogous lineage commitment may take place in the persistent presence of antigen (*35, 36*). We observed the emergence of CX3CR1^+^ cells in the subacute phase even in the absence of TCF-1^+^ cells or CD4 T cell help. Recent studies have described a similar population and concluded that CX3CR1^+^ cells constitute a transitional population downstream of TCF-1^+^ exhausted cells using adoptive transfer (*37*). It is likely that TCF-1^+^ cells retain potential to give rise to CX3CR1^+^ cells as they generate CX3CR1^+^ cells and terminally differentiated cells upon adoptive transfer or PD-1 blockade. However, the conversion from TCF-1^+^ cells to CX3CR1^+^ cells may only slowly occur during the chronic phase of antiviral response or in the tumor microenvironment without PD-1 blockade or adoptive transfer, given that we observe persistence of fate-mapped cells and active proliferation. Together with the slow conversion of *Cx3cr1* fate-mapped cells to TIM3^+^ CX3CR1^−^ cells during the chronic phase, which was also reported by another recent paper (*38*), CX3CR1^+^ cells could constitute an independent lineage that is sustainable by at least limited self-renewal (*38*).

Alternatively, it is possible that CX3CR1^+^ cells require input from TCF-1^+^ cells, albeit at a slow rate, for long-term persistence. Such hierarchical dynamics may be similar to the models regarding the mobilization of hematopoietic stem and progenitor cells, in which hematopoietic homeostasis is maintained predominantly by progenitors with limited self-renewing capability while stem cells are necessary for long-term hematopoiesis but only rarely give rise to these progenitors (*14, 15*). Analogously, TCF-1^+^ cells act as reserve cells that can be called upon in very specific contexts such as checkpoint blockade, but likely do not contribute actively to CD8 T cell responses. Instead, TIM3^−^ CX3CR1^+^ cells, which are likely maintained by the IL-21-T-bet axis, serve as an intermediate progenitor population that constantly gives rise to CX3CR1^+^ TIM3^+^ cells as effectors. Thus, there is a hierarchy of CD8 T cells beginning with TCF-1^+^, and ending with TIM3^+^ CX3CR1^+^ or TIM3^+^ CX3CR1^−^ cells potentially through separate developmental pathways. This hierarchy could be essential for durable responses as it would prevent depletion of a long-term precursor pool during long-lasting infections, potentially resulting in clonal depletion.

An important takeaway of our work is the remarkable persistence of CD8 T cells that fall into the exhausted state during active infection. Specifically, while stem-like TCF-1^+^ cells have been demonstrated to be long-lived, this phenotype is also the case for CX3CR1^+^ cells. A substantial fraction of TCF-1^+^ memory CD8 T cells found after viral clearance are derived from CX3CR1^+^ cells. Prior studies have utilized *Tcf7*-deficient mice to demonstrate a requirement for the stem-like population for persistence under competitive conditions, however, TIM3^−^ CX3CR1^+^ cells do express a low amount of TCF1, providing an alternative explanation for overall cell loss. In addition, mice used for the transfer studies lack significant CD4 T cell responses, which likely secondarily results in loss of CX3CR1 cells (*8–10*). New tools that allow for fate mapping directly from the TCF-1^+^ pool will help us to understand the extent of their dynamics and in different contexts.

T cell fates following infection have been defined as the formation of effector cells and memory precursor cells in acute infection from naive cells. However, recent data indicate that this process is not necessarily unidirectional with KLRG1^+^ short-lived effector cells able to de-differentiate into memory cells with considerable proliferative capacity (*39*). In addition, deletion of the effector-promoting transcription factor Id2 results in the default acquisition of memory phenotypes (*40*). Here we demonstrate a strikingly similar phenomenon even in exhausted CD8 T cells, with conversion of CX3CR1^+^ BLIMP-1^+^ TCF-1^lo/–^ cells to CX3CR1^+^ TCF-1^+^ cells after the infection is resolved. This result indicates cells other than TCF-1^hi^ stem- or central memory-like cells retain the capacity to re-express TCF-1 under certain conditions, with undefined microenvironmental or intrinsic factors promoting this transition, possibly mediated by epigenetic reprogramming as reported in acute response (*41*). We expect to observe an equivalent type of memory T cell following cancer elimination, which may be important to prevent clinically detectable relapse. It remains to be determined how this memory cell type differs from CX3CR1^−^ memory cells though the acquisition of imprinted identity during the effector phase. This knowledge will potentially enhance long-term immunity against chronic viral infection or cancers.

## Materials and Methods

### Mice and Infection

Male C57BL/6N and B6-CD45.1 mice were purchased from Charles River Laboratory. *Prdm1-EYFP,* CD8 (*E8I)-Cre, Tbx21^F^, Eomes^F^, Bcl6^F^, Il21r, Cx3cr1^CreER^, Rosa26^LSL-tdT^–* mice were originally obtained from The Jackson Laboratory. All mice were housed in a specific pathogen-free facility at Washington University in St. Louis, and were used for infection at 8 to 10 weeks of age unless stated otherwise. All experiments were performed according to a protocol approved by Washington University’s Institutional Animal Care and Use Committee (IACUC). Stocks of LCMV were made by propagating virus by infection of BHK (baby hamster kidney) cells, followed by titering of culture supernatants by focus forming assay on Vero (African green monkey kidney) cells. For LCMV infection, mice were infected with 2 x 10^5^ plaque-forming units (PFU) of LCMV-Arm strain via the intraperitoneal route or 2 x 10^6^ (PFU) of LCMV-c13 by intravenous injection. For the quantification of plasma viral load, RNA was extracted from 10 μl of plasma using Trizol (Life Technologies). Before RNA extraction a spike-in of exogenous control ERCC RNA was carried out and used to normalized viral loads following qPCR as described previously (*31, 42*).

### Bulk RNA-Seq

To examine expression of different CD8 T cell subsets we utilized *Prdm1-EYFP* mice to separate TCF-1^+^ (YFP^−^) and TCF-1^−^ (YFP^+^) cells in the LCMV-specific CD8 T cells as previously described (*9, 43*). CD8^+^ T cells were first enriched from splenocytes using the Dynabeads FlowComp Mouse CD8 T cell kit followed by surface staining. We then sorted *Prdm1-EYFP*^−^ TIM3^−^ CX3CR1^−^ cells, which corresponded to TCF-1^+^ stem-like CD8 T cells, and divided *Prdm1*-EYFP^+^ cells into CX3CR1^+^ TIM3^−^, CX3CR1^+^ TIM3^+^, and CX3CR1^−^ TIM3^+^ populations from splenocytes 16 days after infection with LCMV-c13.

Total RNA was extracted from ∼20-50K sorted cells using the RNA XS Kit (Macherey Nagel) according to the manufacturer’s instructions. cDNA synthesis and amplification were performed with Next Ultra RNA Library Preparation Kit (NEB). Libraries were sequenced on a HiSeq3000 (Illumina) in single-read mode, with a read length of 50 nucleotides producing ∼25 million reads per sample. Sequencing reads were mapped to NCBI mm9 using STAR with default parameters, and mapping rates were higher than 90%. Transcripts with 4 TPM at least in one sample were initially filtered, followed by principal component analysis, unsupervised clustering, and profiling of subpopulation-specific gene expression using Phantasus and limma packages (V.15.1, https://artyomovlab.wustl.edu/phantasus/). The raw data will be deposited at NCBI GEO upon acceptance of the manuscript.

### Single-cell RNA-seq

Single-cell RNA-seq libraries were prepared using the 10X Single Cell Immune Profiling Solution Kit (v1 Chemistry), according to the manufacturer’s instructions. Briefly, FACS sorted cells were washed once with PBS + 0.04% BSA. Following reverse transcription and cell barcoding in droplets, emulsions were broken and cDNA purified using Dynabeads MyOne SILANE followed by PCR amplification (98C for 45 sec; 14 cycles of 98C for 20 sec, 67C for 30 sec, 72C for 1 min; 72C for 1 min). For gene expression library construction, 50 ng of amplified cDNA was fragmented and end-repaired, double-sided size selected with SPRIselect beads, PCR amplified with sample indexing primers (98C for 45 sec; 14 cycles of 98C for 20 sec, 54C for 30 sec, 72C for 20 sec; 72C for 1 min), and double-sided size selected with SPRIselect beads. The prepared single-cell RNA libraries were sequenced on an Illumina HiSeq 4000 to a minimum sequencing depth of 25,000 reads/cell using the read lengths 28bp Read1, 8bp i7 Index, 91bp Read2. Single-cell RNA-seq reads were aligned to the mm10 reference genome and quantified using cellranger count (10X Genomics, version 3.1.0). Filtered gene-barcodes matrices containing only barcodes with UMI counts passing threshold for cell detection were used for further analysis.

Additional analysis was performed using Seurat (version 3.1.2) (*44*). Cells with less than 200 genes detected or greater than 5% mitochondrial RNA content were excluded from analysis. For clustering, raw UMI counts were log normalized and variable genes identified based on a variance stabilizing transformation. We assigned scores for S and G2/M cell cycle phase based on previously defined gene sets (*45*) using the CellCycleScoring function. Scaled z-scores for each gene were calculated using the ScaleData function and regressed against the S phase score and G2/M phase score to reduce clustering based on cell cycle. Scaled z-scores for variable genes were used as input into PCA. Clusters were identified using shared nearest neighbor (SNN) based clustering based on the first 10 PCs with k = 15 and resolution = 0.23. The same principal components were used to generate the UMAP projections (*46, 47*), which were generated with a minimum distance of 0.1 and 30 neighbors. Expression of selected genes was plotted using log normalized gene expression values.

### Treatments

For depletion of CD4 T cells, 200 micrograms of anti-CD4 (GK1.5, Leinco) was injected on −1 and +1 dpi. For administration of Tamoxifen (Sigma), 10 mg/mL solutions were prepared in Corn Oil (Sigma) and 1mg gavaged orally.

### Cell Preparation, Cell Staining, and Flow Cytometry

Single-cell suspensions of splenocytes were prepared by manual disruption with frosted glass slides. Lungs were minced with scissors and digested with Collagenase D (Sigma) and DNase I (Sigma) with agitation for 1 hour at 37C followed by enrichment of lymphocytes by a 40/70 Percoll gradient. Absolute live cell counts were determined by Trypan-blue exclusion using Vi-Cell (Beckman Coulter). Tetramer staining was performed using iTag-PE and APC LCMV gp33-44 and gp276-284 (MBL international). The following monoclonal antibodies were purchased from Biolegend unless otherwise indicated: FITC-conjugated anti-CD45.2 (104); PerCP-Cy5.5 conjugated anti-CD8β (YTS156.7), anti-CD4 (GK1.5), anti-Ki-67 (16A8) PerCP-eFlour710-conjugate anti-Eomes (eBioscience, Dan11mag), APC-conjugated CD366 (TIM-3, RMT3-23), anti-TNF-α (MP6-XT22), anti-granzyme B (QA16A02), anti-T-bet (4B1), anti-CX3CR1 (SA011F11), Alexa 700-conjugated anti-CD44 (IM7), anti-CD45.2 (104), BV421-conjugated anti-TIM3 (RMT3-23), BV605-conjugated CD4 (GK1.5), anti-CD45.1 (A20), anti-CX3CR1(SA011F11), BV711-conjugated anti-CD4 (RM4-5), PE-conjugated anti-IFN-γ (XMG1.2), anti-PE-Cy7-conjugated anti-PD-1 (29F.1A12) PE-Dazzle594-conjugated anti-B220 (RA3-6B2) and BUV395-conjugated anti-CD8 (53-6.7, BD) Anti-TCF-1 antibody (Cell Signaling, C63D9) and detected by Alexa 488-conjugated Donkey anti-rabbit polyclonal IgG (Thermo Fisher, Cat. #R37118). Staining for transcription factors was performed using the Foxp3 staining kit (eBioscience) according to the manufacturer’s instructions.

For intracellular cytokine staining, splenocytes were cultured in RPMI-1640 supplemented with 10% fetal bovine serum in the presence of 1 ug/ml of LCMV-gp peptide (Genscript) and 5 ug/ml of Brefeldin A (Biolegend) for 4 hours. Cells were stained for surface makers and then subject to LIVE/DEAD Aqua staining (Thermofisher) for 30 minutes at 4°C before being fixed with 4% PFA for 10 minutes at R. Cells were then washed twice with 0.03% saponin in 2% FBS/PBS before being stained with the indicated antibodies in 0.3% Saponin in 2% FBS/PBS for 20 min at 4°C.

BrdU incorporation assay was performed using the APC BrdU Flow Kit (BD Biosciences) according to the manufacturer’s instruction with one-time intraperitoneal injection of 1 mg of BrdU followed by tissue collection 12 hours later.

Stained samples were analyzed with FACS LSR Fortessa, X20 (BD), or Symphony A3 or sorted on Aria II or III. Data were processed with FlowJo Software (FlowJo. LLC).

### Tumor Inoculation and Analysis

EG7 (ATCC) and MC38 (generous gift of Arlene Sharpe) cells were grown in RPMI and DMEM, respectively supplemented with 10% FBS (Gibco), GlutaMAX (Gibco), and b-Mercaptoethanol. Mice were shaved one day prior to subcutaneous inoculation of 10^6^ cells. For TIL analysis tumors were excised, minced, and digested with Collagenase B (Sigma), and DNase I for 30 minutes at 37°C to prepare single cell suspension.

### Statistical analysis

*P-*values were calculated with an unpaired two-tailed Student’s *t*-test or Mann-Whitney U-test for two-group comparisons and by one-way ANOVA for multi-group comparisons with the Tukey post hoc test using Prism 8 software. \**P* < 0.05; ** *P* < 0.01; *** *P* < 0.001; **** *P* < 0.0001.

## Supplementary Materials

**Supplementary Fig. 1.**
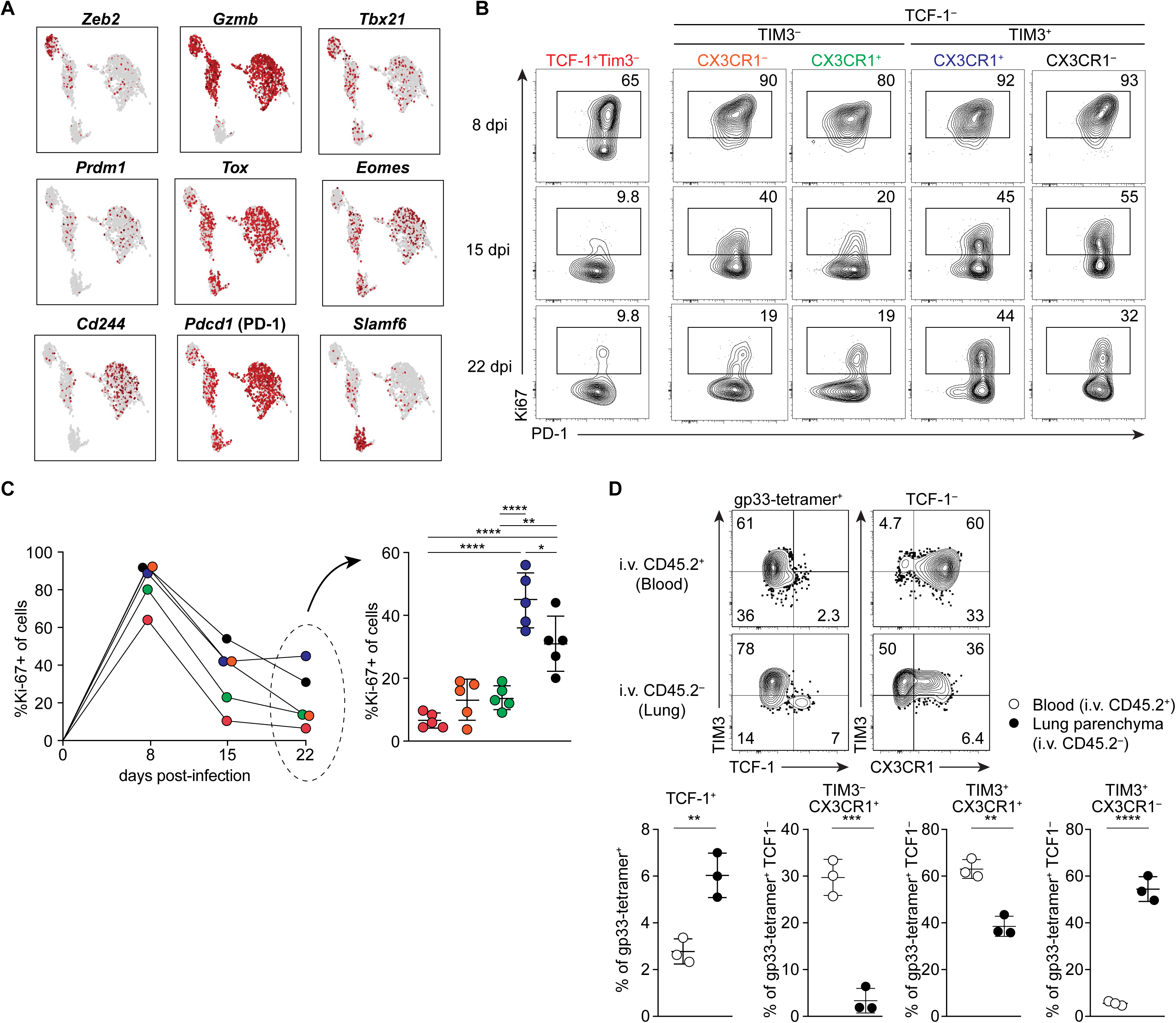
Characterization of CX3CR1 expressing CD8 T cells in infection and tumors. **A.** UMAP analysis of scRNA-seq data of gp-33-specific CD8 T cells on 21 dpi with LCMV-c13 showing expression of indicated genes. **B, C.** Expression of PD-1 and Ki-67 in subpopulations of gp33-specific CD8 T cells defined by TIM3 and CX3CR1 in LCMV-c13-infected C57BL/6 mice at different time points after infection. Data are representative of two experiments with n=5 mice. **D.** Flow cytometry analyzing expression of TIM3, CX3CR1, and TCF-1 by LCMV gp33-specific CD8 T cells in lungs of mice that were intravenously injected with 3 ug of CD45.2 three minutes before euthanasia. Data are representative of 2 independent experiments with n=3 mice each. Each dot in the graphs indicates an individual mouse that was examined, and data are shown by mean ± SD. Statistical differences were tested using one-way ANOVA with a Tukey post-hoc test (C) and student’s t-test (D).

**Supplementary Fig. 2.**
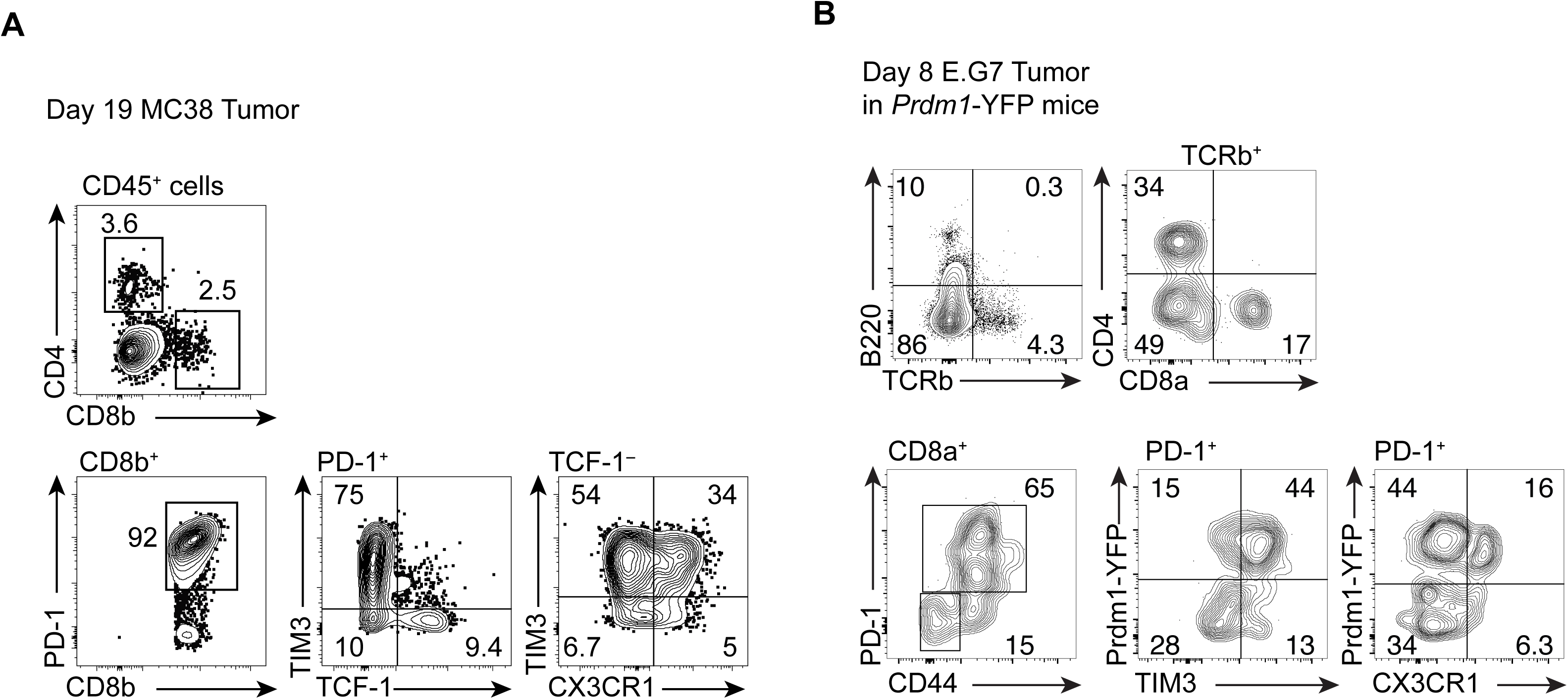
Analysis of CX3CR1 Expression in TIL. **A, B.** Expression of TCF-1, TIM3 and CX3CR1 in PD-1^+^ CD8 T cells infiltrating in the MC38 tumors (A) 19 days and E.G7 tumors (B) 8 days after subcutaneous inoculation. Data are representative of 3 independent experiments with n>3 mice each.

**Supplementary Fig. 3.**
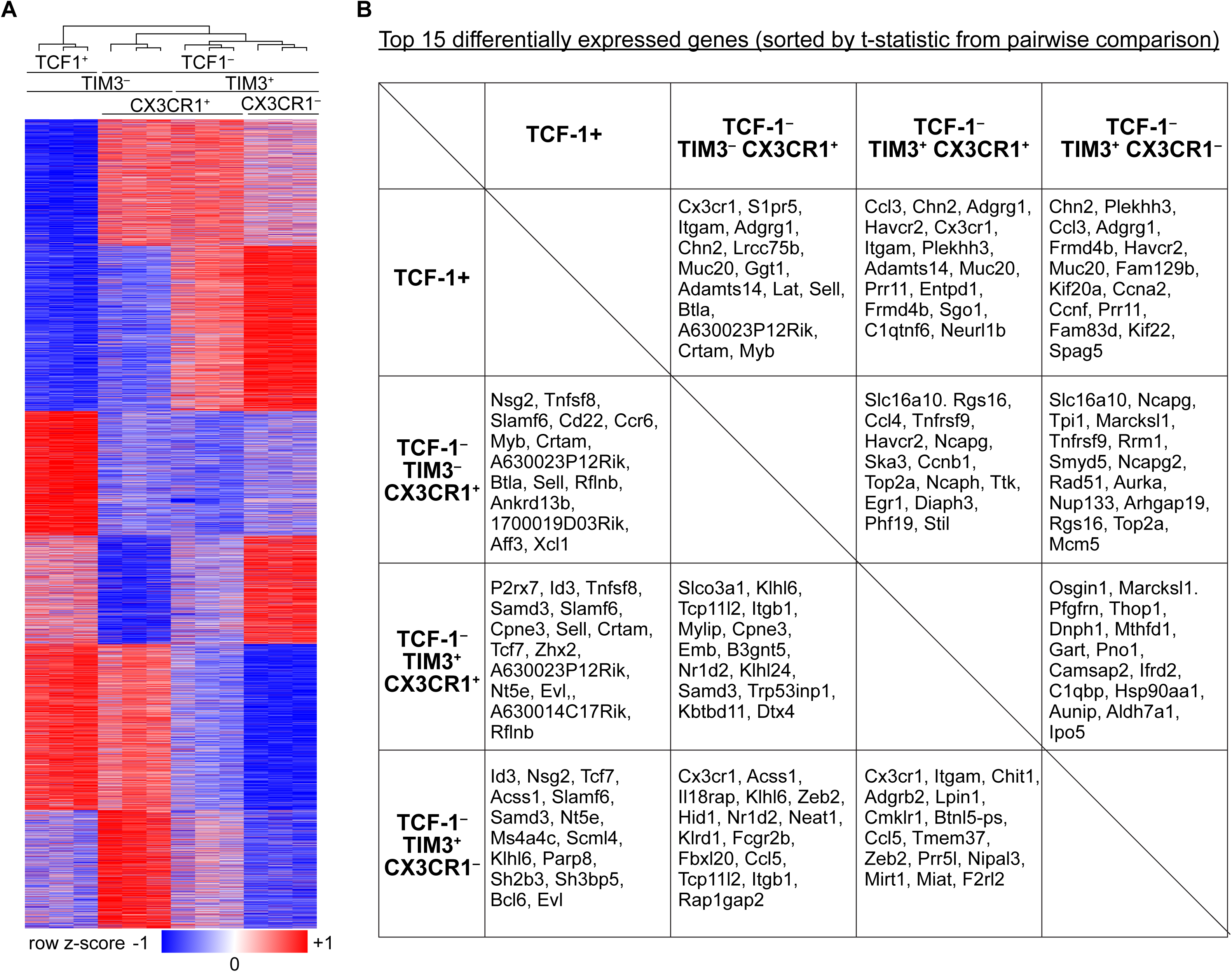
Profiling of differentially expressed genes between distinct subpopulations of PD-1^+^ CD8 T cells following LCMV-c13 infection. **A.** Heatmap showing K-mean clustering of genes differentially expressed between distinct subpopulations on 16 dpi. Gene expression levels are shown as z-score normalized to each row mean. **B.** Table showing top 15 differentially expressed genes for each pairwise between two indicated populations, as determined by significance based on t-statistics. Genes shown are more highly expressed in the population listed horizontally on the top row compared to one listed vertically on the left side.

**Supplementary Fig. 4.**
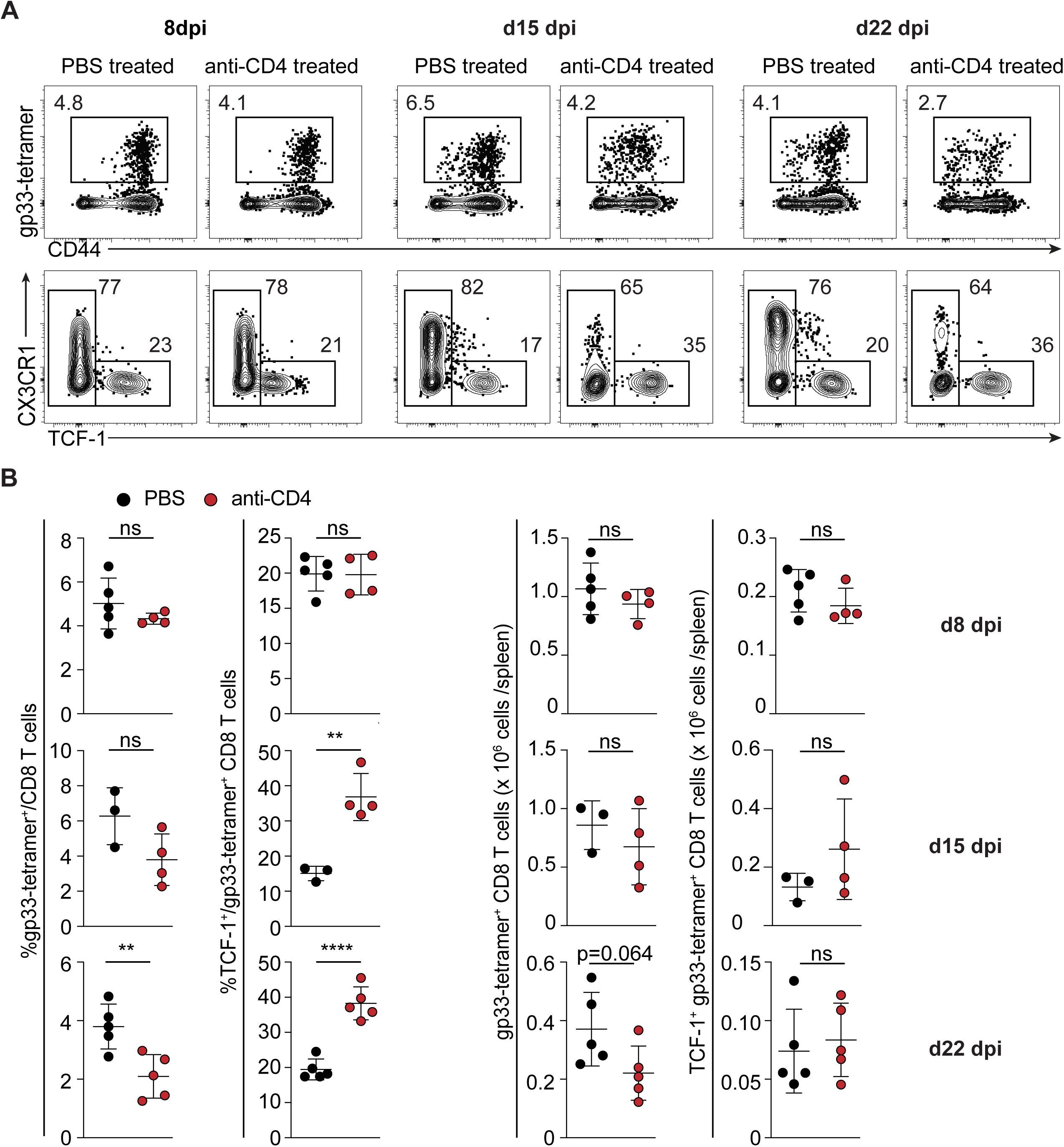
Time-course of CD4 T cell dependent CD8^+^ T cell response during LCMV-c13 infection. **A, B.** Flow Cytometry showing expression of CD44 and H-2Db(gp33-41)-specific TCR of total CD8 T cells as well as expression of TCF-1 and TIM3 in gp33-specific CD8 T cells in the spleen of control and CD4 T cell-depleted mice on the indicated time points after infection. (A) shows representative plots from 3-5 mice and data from replicates are shown in (B) with mean ± SD.

**Supplementary Fig. 5.**
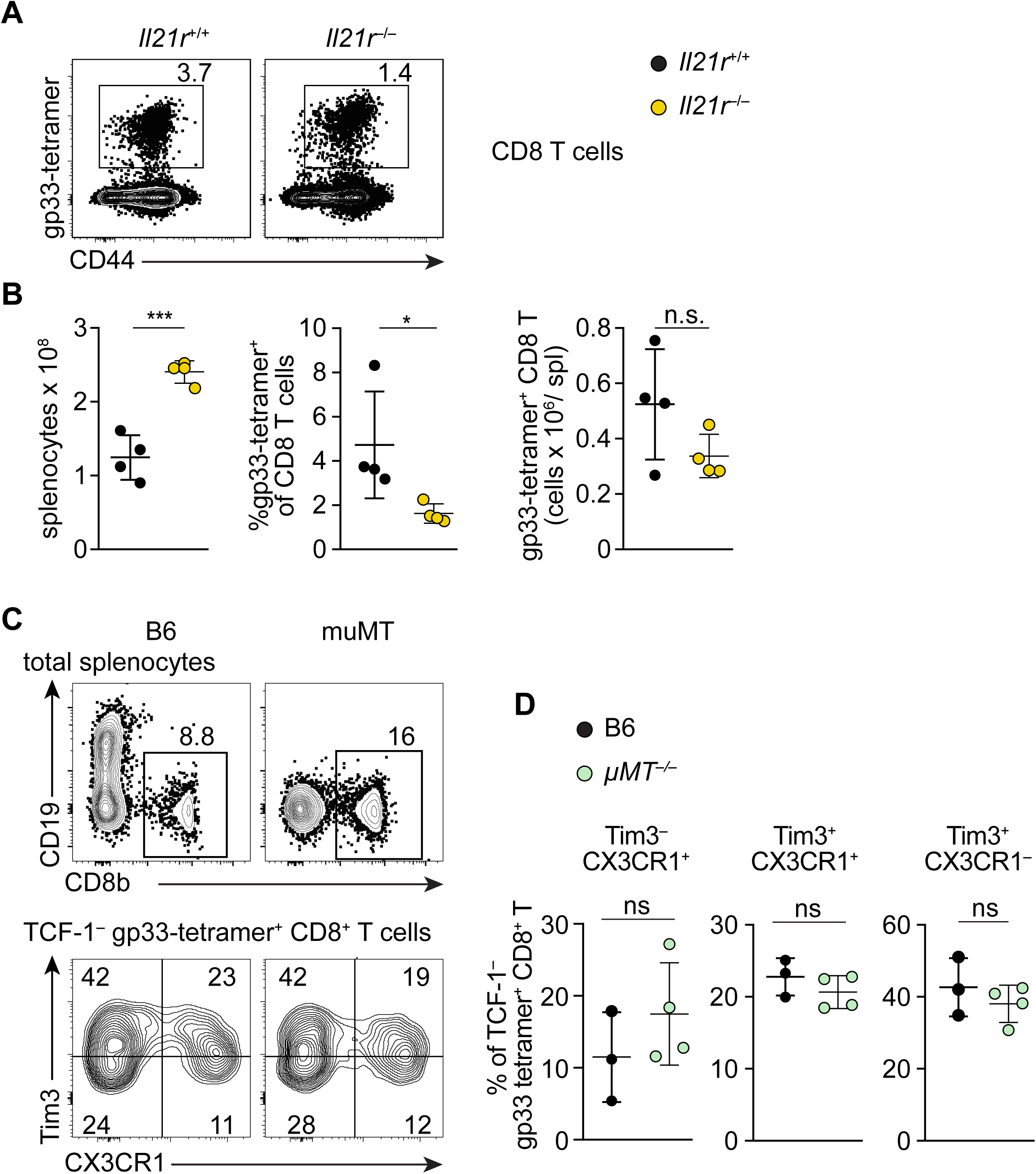
IL-21 signaling independent of germinal center response is necessary for the maintenance of CX3CR1+ PD-1+ CD8 T cells in LCMV-c13-infected mice. **A, B.** Flow cytometry showing expression of LCMV-gp33-specific TCR and CD44 in CD8 T cells in *Il21r*^+/+^ and *Il21r*^−/–^ mice on 35 dpi. Data are representative of two independent experiments with n>3 mice in each experiment per genotype. **C, D.** Flow cytometry showing expression of TIM3 and CX3CR1 in TCF-1^−^ gp33-specific CD8 T cells in muMT mice. Representative plots shown in (C) with data from replicates in (D) with mean ± SD.

**Supplementary Fig. 6.**
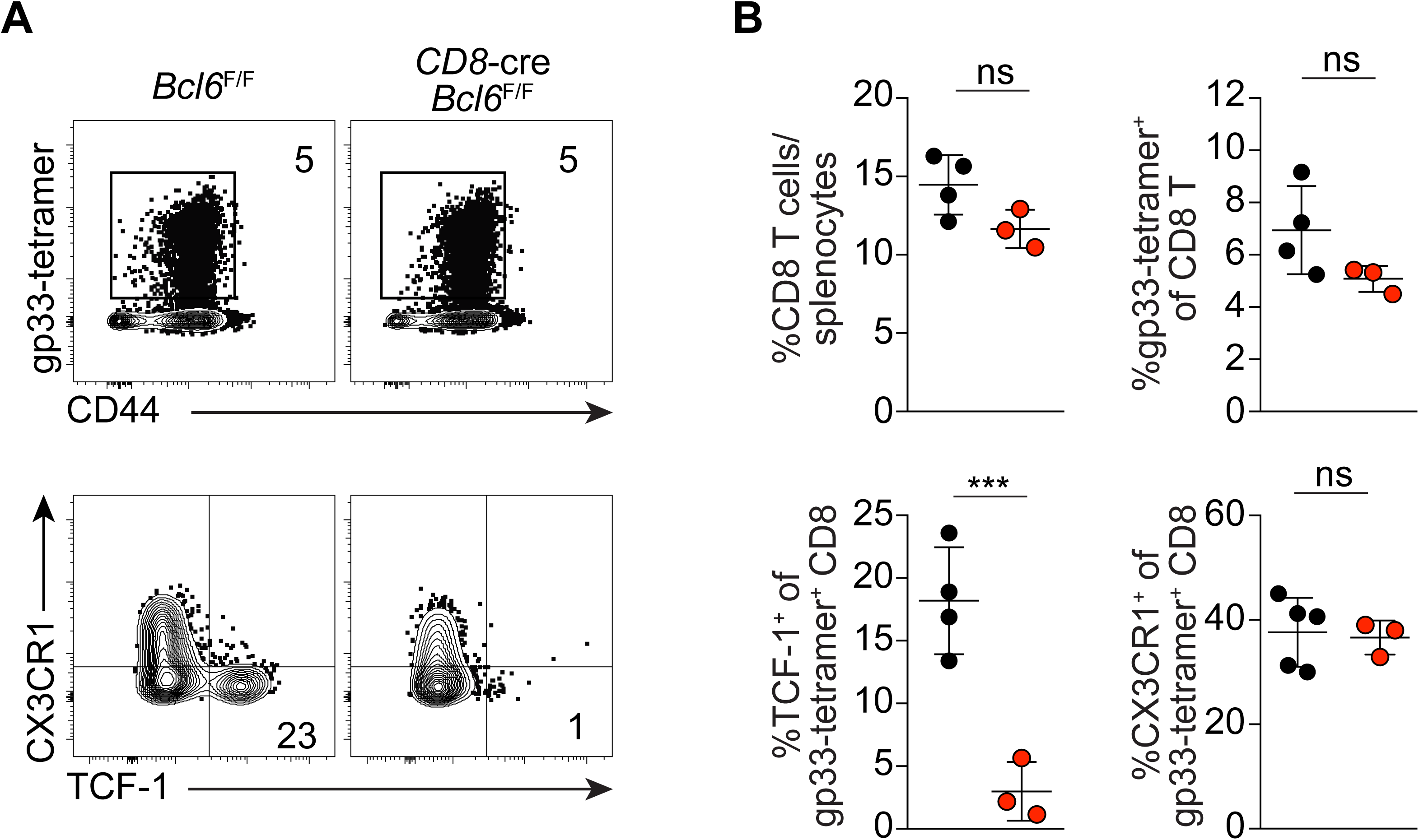
Bcl6 is required for the formation of TCF-1^+^ stem-like CD8 T cells but not for CX3CR1^+^ cells in LCMV-c13 infection. Flow cytometry data showing TCF-1 and CX3CR1 in gp33-specific CD8 T cells on 8 dpi (A). Data from replicates are shown in (B) with mean ± SD.

**Supplementary Fig. 7.**
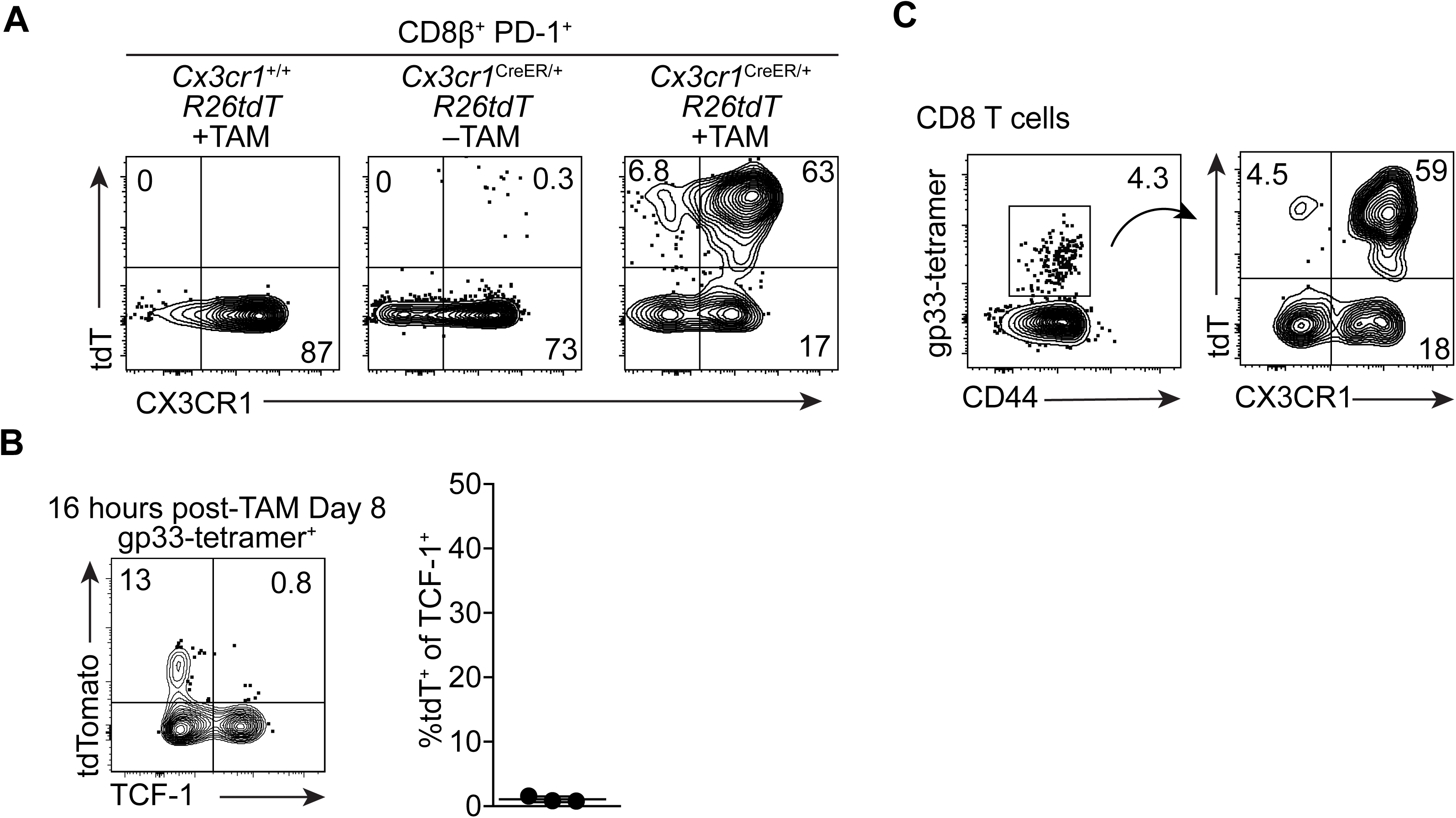
Characterization of *Cx3cr1*^creERT2/+^ *ROSA26*^LSL-tdT/+^ mice for *in vivo* labelling. **A.** Flow cytometry plots showing expression of tdT and CX3CR1 in the PBMC of mice on 10 dpi in the indicated genotypes of mice following oral administration of Tamoxifen on 8 and 9 dpi. **B.** Flow cytometry plots showing expression of tdT and TCF-1 in gp33-specific CD8 T cells 16 hours following tamoxifen administration on 8 dpi. Data from replicates are shown in the right panel. Data are representative of 2 independent experiments with at least 3 mice each. **C.** Flow cytometry plots showing expression of tdT and CX3CR1 in gp33-specific CD8 T cells 24 hours following tamoxifen administration on 22 dpi. Data are representative of 2 independent experiments with at least 3 mice each.

## Acknowledgements

We thank M. Colonna and M. Cella for LCMV stocks, G. Randolph for *Cx3cr1*-creERT2 mice, Arlene Sharpe for MC38 cells, C. Fuji for the maintenance of the mouse colony, and C-S. Hsieh for discussion and critical reading of the manuscript. This study was supported by NIH grants R01AI130152-01A1 and R03AI139875-01 (to T.E.), T32HL007317 (to S.R.), and T32GM007200 (to S.R. and D.J.V.). T.E. is a Scholar of the Leukemia and Lymphoma Society (www.lls.org).

## Author Contributions

SR and TE designed the study. SR, YX, BD, KEY, ET, EB, DJV and TE conducted experiments. SR, YX, ATS and TE interpreted results. SR and TE wrote the manuscript with comments from all authors.

## Data Availability

Bulk RNA-seq data are deposited to NCBI Gene Expression Omnibus under the accession number GSE145910. Data will be made publicly accessible upon acceptance of the manuscript.

